# Simultaneous single cell imaging of calcium signal dynamics in breast cancer and neural cells reveals communication in a model of brain metastasis

**DOI:** 10.1101/2025.07.23.666251

**Authors:** Silke B Chalmers, Alice H L Bong, Francisco Sadras, Mélanie Robitaille, Jodi Saunus, Sunil R Lakhani, Sarah J Roberts-Thomson, Gregory R Monteith

## Abstract

The brain provides a unique metastatic microenvironment for breast cancer cells, where calcium signaling dynamics play critical roles in both cancer cell behavior and normal brain function. Calcium signal-mediated communication between breast cancer and neural cells has not yet been demonstrated through selective activation of breast cancer cells at the single cell level. To address this, we combined neural matrices of differentiated human neural progenitor cells with breast cancer cells expressing spectrally distinct genetically encoded calcium indicators. Specific activation of breast cancer cells increased calcium signaling activity in neural matrices with distinct temporal and spatial characteristics. Neural matrices also remodeled the expression of calcium-sensitive transcription factor SOX2 in a manner dependent on proximity to breast cancer cells. This work is the first simultaneous single-cell assessment of calcium signaling dynamics between breast cancer cells and neural cells modelling interactions within the brain metastatic niche.

## Introduction

The brain is unique among sites of breast cancer metastases. The coalescence of limited treatment options, high disease burden and low survival rates culminate in uniquely poor patient prognoses and a significant clinical challenge^1–5^. While treatments harnessing the tumor microenvironment have improved outcomes in the primary breast cancer setting, comparable options for brain metastases are lagging^1,4,6–9^. A contributing factor to this delay is the distinct cellular landscape of the brain, which consists of neurons and glial cells such as astrocytes and oligodendrocytes^7^. Intracranial metastases invoke specific and complex adaptations in both invading neoplastic cells and the surrounding microenvironment in comparison to other sites of metastases^1,10–18^. The emergence of these adaptations is contingent on ongoing communication between metastatic breast cancer cells and the brain microenvironment, enabling their survival and growth in the unfamiliar neural landscape. Defining the suite of possible communication mechanisms between breast cancer cells and neural cells has the potential to illuminate novel avenues for therapeutic intervention. One pathway which has been gaining interest as a potential method of communication between breast cancer cells and the brain microenvironment is calcium ion (Ca^2+^) signaling.

Ca^2+^ is a ubiquitous second messenger essential for a myriad of cellular and tissue level processes^19–21^. Dysregulation of Ca^2+^ signaling is a feature of many cancers, and therapies targeting this dysregulation have begun to enter clinical trials due to their effects on cancer cell proliferation^22–25^. There has been comparatively less exploration into the role that Ca^2+^ signaling may play in the context of metastatic progression, particularly in the brain^23^. Ca^2+^ signaling is a critical component of neurological function and recent work has demonstrated that Ca^2+^ communication between neoplastic and neural cells is an essential dependency for glioma survival, a vulnerability that could be targeted to reduce tumor growth^26–28^. Whether metastatic breast cancers could employ a similar survival strategy within the brain microenvironment was largely unexplored until Zeng *et al*. identified the presence of “pseudo-tripartite synapses” between breast cancer cells and glutamatergic neurons via electron microscopy^29^. Such structures were proposed to drive metastatic breast cancer survival due to Ca^2+^ communication via N-methyl-D-aspartate receptor signaling. However, the functionality of these synapses was not proven as live cell signaling assays were not conducted in co-cultures. Further exploration of the potential role of Ca^2+^ communication between breast cancer and neuronal cells have occured^30,31^. However, simultaneous assessment of Ca^2+^ signaling dynamics in breast cancer and neural cells, the essential direct evidence of Ca^2+^ communication between metastases and the microenvironment, has still yet to be conducted. To address this gap, we generated a breast cancer and neural microenvironment model using human neural progenitor cell line ReNcell VM and triple negative breast cancer cells MDA-MB-468, with expression of spectrally distinct genetically encoded Ca^2+^ indicators, to characterize Ca^2+^ signaling dynamics between breast cancer cells and neural cells through selective activation of Ca^2+^ influx in breast cancer cells. This model demonstrated that breast cancer cells can initiate localized Ca^2+^ signaling in human neural cells, and reveals a previously undocumented example of breast cancer–neural cell interaction.

## Results and Discussion

### Human neural progenitor cell line ReNcell VM differentiates into electro-physically responsive microenvironments that support breast cancer survival and growth

Studies exploring breast cancer in brain microenvironments have been conducted in in vitro models consisting of a single neural cell type, such as primary rodent astrocytes or microglia cells or with ex vivo studies involving whole brain slices or organotypic models lacking single cell resolution of Ca^2+^ signaling or specific activation of Ca^2+^ influx in breast cancer cells^17,32–36^. We utilized the immortalized human neural progenitor cell line, ReNcell VM to model the brain milieu and to allow high throughput single cell resolution of Ca^2+^ signaling^37^. Human neural progenitor cells are regularly maintained in a proliferative, stem-like state, but upon mitogen withdrawal will undergo a differentiation process resulting in neural matrices consisting primarily of astrocytes and neurons^37–41^. Indeed, withdrawal of mitogens from proliferative ReNcell VM resulted in upregulated mRNA expression of astrocytic marker glial fibrillary acidic protein (GFAP) and neuronal marker MAP2 after 4 days (Fig 1a). Differentiated ReNcell VM developed into interconnected neural matrices with spindle-like morphology that stained positively for GFAP and neuronal marker β3-tubulin (Fig 1b). Automated analysis of cell proportions found the matrices consisted of 63 – 77% GFAP positive cells, and 6.5 - 10% β3-tubulin positive cells (Fig 1c). These proportions reflect the ratio of neuronal to non-neuronal cell types seen in the ventral mesencephalon area of the midbrain, the region from which the ReNcell VM line was originally isolated^37,42^. They are also consistent with previous reports that 4% of ReNcell VM cells within matrices are β3-tubulin positive after 4 days of differentiation, as assessed by fluorescence-activated cell sorting^43^. Following morphological and expression assessments, we sought to determine whether differentiated ReNcell VM matrices were electro-physiologically responsive. To do this, we stably expressed the genetically encoded Ca^2+^ indicator jRCaMP1b in the ReNcell VM model and assessed changes in intracellular Ca^2+^ levels via automated epi-fluorescence microscopy (Fig 1d)^44^. Development of spontaneous Ca^2+^ activity is typically observed in the shift from neural progenitor cell to a mature neural state, and assessment of differentiated ReNcell VM matrices identified frequent spontaneous Ca^2+^ activity (Fig 1d)^45^. Moreover, differentiated ReNcell VM matrices were highly responsive to addition of the muscarinic agonist carbachol, an agent commonly utilized for its ability to elicit Ca^2+^ mobilisation in neural cultures including primary astrocytes and neurons (Fig 1d)^46–48^. Collectively these results indicated that differentiated ReNcell VM neural matrices recapitulate key aspects of the brain milieu, including cellular composition and electrophysiological properties, and, as such, we sought to confirm whether this model would be suitable for co-culture studies with human breast cancer cells.

**Fig. 1:**
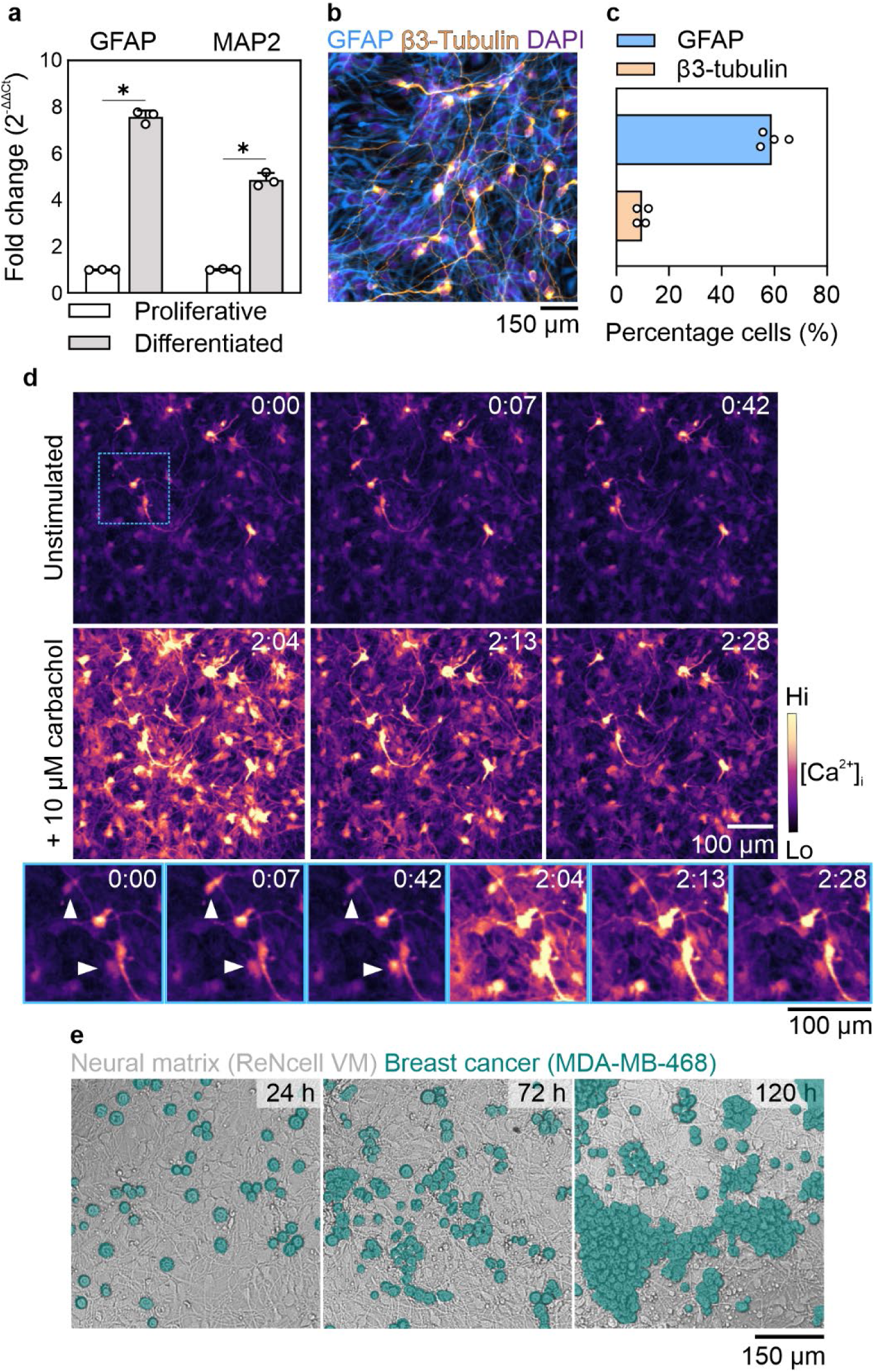
Human neural progenitor cell line, ReNcell VM, as an in vitro model of the brain microenvironment. **a** ReNcell VM induced to undergo differentiation increase expression of astrocytic marker GFAP and neuronal marker MAP2. Graph shows mean + SD, n=3 independent experiments conducted in duplicate, unpaired two-tailed t-test, \**p* < 0.05. **b** Representative immunofluorescent staining of astrocytic marker GFAP (blue), neuronal marker β3-tubulin (orange) and nuclear marker DAPI (purple) in differentiated ReNcell VM neural matrices **c** Proportion of cells within differentiated ReNcell VM neural matrices positively stained for GFAP or β3-tubulin. Graph shows mean. Data from four microplate wells, two independent experiments. **d** Differentiated neural matrices have characteristic neural Ca^2+^ signaling properties, including spontaneous activity (top series, excerpt in blue), and responses to muscarinic agonist carbachol (bottom series). Arrows highlight spontaneously active cells. Timer indicates minutes:seconds. **e** Representative brightfield images of breast cancer cell line MDA-MB-468 surviving and proliferating while co-cultured on differentiated ReNcell VM neural matrices. Breast cancer cells are highlighted in green. Images representative of three independent experiments.

While ReNcell VM are commonly utilized for neurological investigations, to date very few have assessed their utility for co-culture studies^49^. We introduced a triple-negative breast cancer cell line (MDA-MB-468) onto differentiated ReNcell VM neural matrices and maintained these cells in co-culture (Fig 1e, Fig S1). MDA-MB-468 breast cancer cells were not only able to survive within this co-culture, but proliferated in this environment (Fig 1e, Fig S1). While most MDA-MB-468 cells grew in breast cancer cell clusters, examples of single cells could also be observed within the neural matrix even after 120 h of culturing (Fig 1e). ReNcell VM matrices remained intact throughout co-culturing with MDA-MB-468 cells and did not deteriorate as a consequence of breast cancer cell addition and proliferation (Fig 1e).

### Selective stimulation of Ca^2+^ influx in MDA-MB-468 breast cancer cells increases Ca^2+^ activity in co-cultured neural matrix

The interplay between breast cancer and metastatic microenvironment plays a pivotal role in the survival of invading neoplasms to foreign sites^2,13,14,33,50^. Ca^2+^ signaling has been proposed as an important communication avenue in brain metastasis, however, to date, no studies have simultaneously assessed Ca^2+^ signaling between breast cancer and neural cells in a live cell model^29–31,50^. We hypothesized that communication of a Ca^2+^ signal between neural and breast cancer cells could be assessed via the administration of a pharmacological activator able to selectively increase cytosolic Ca^2+^ in breast cancer cells. To this end, we screened differentiated ReNcell VM matrices and MDA-MB-468 breast cancer cells for unique expression of Ca^2+^ channels and receptors with selective pharmacological activators. We have previously reported that certain breast cancer cell lines, including MDA-MB-468 cells, express high levels of the plasma membrane resident Ca^2+^ permeable ion channel transient receptor potential vanilloid 4 (TRPV4)^51^. This channel has a highly selective pharmacological activator, GSK1016790A which induces robust and sustained Ca^2+^ influx in MDA-MB-468 breast cancer cells at nanomolar concentrations^51^. Assessment of mRNA expression of TRPV4 in differentiating ReNcell VM demonstrated that expression of this channel was undetectable after 7 days of differentiation (Fig 2a). As expected, stimulation of differentiated ReNcell VM monolayers with TRPV4 activator GSK1016790A did not cause a detectable increase in Ca^2+^ influx (Fig 2b&c). Having identified an appropriate Ca^2+^ influx inducing agent, we assessed Ca^2+^ signalling in neural co-culture after selective activation of Ca^2+^ influx in breast cancer cells.

**Fig 2:**
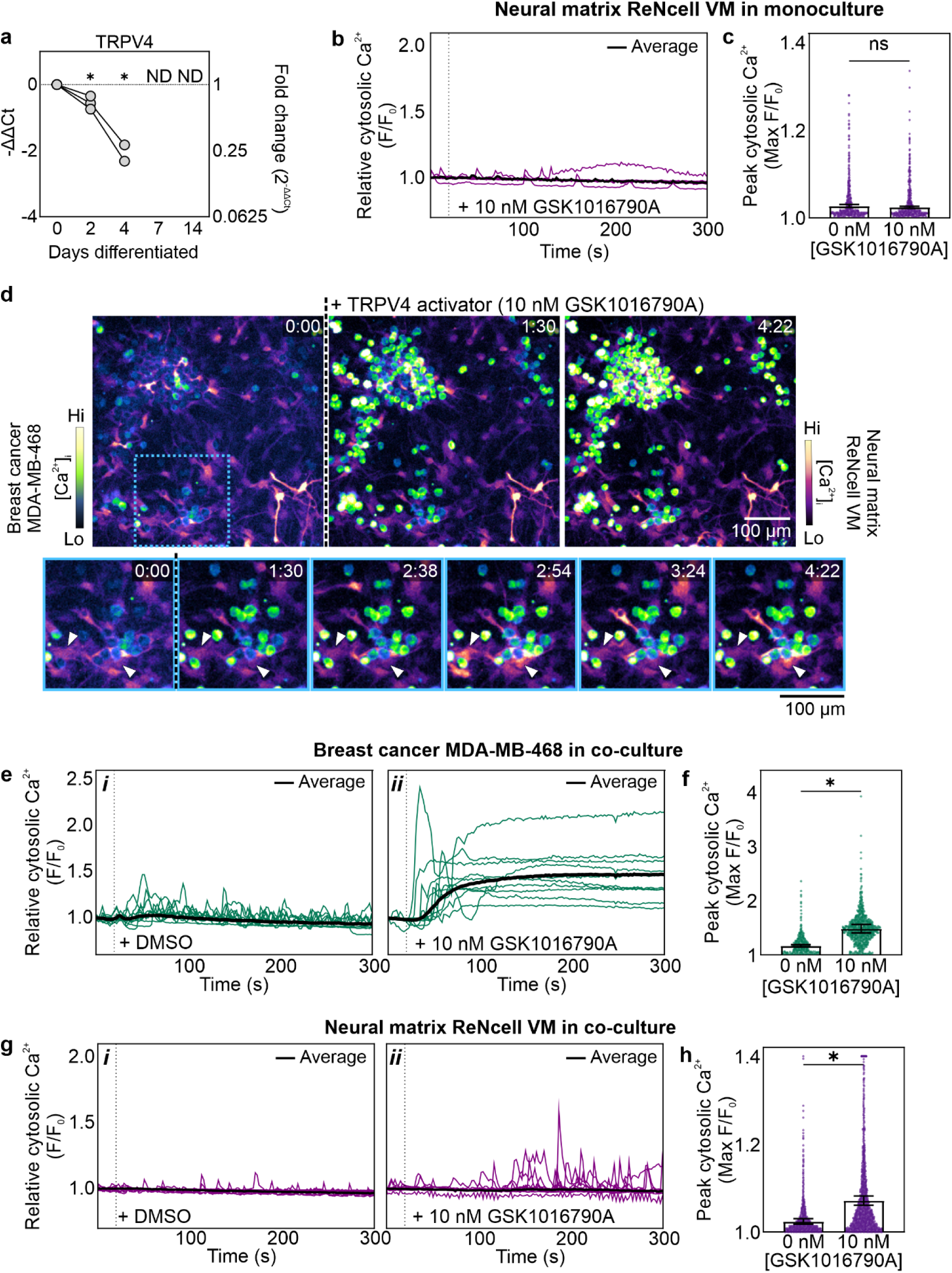
Stimulated Ca^2+^ signaling in breast cancer cells increases Ca^2+^ activity in the neural microenvironment. **a** TRPV4 mRNA expression in ReNcell VM matrices following induction of differentiation. Graph depicts three independent experiments conducted in duplicate. Results were analysed via one-way ANOVA with Tukey’s multiple comparison’s test. *Indicates significance in comparison to 0 d differentiated (\**p* < 0.05). ND; not detected. **b** Ca^2+^ responses in differentiated ReNcell VM neural matrices following administration of TRPV4 activator (10 nM GSK1016790A, time of addition indicated by dashed line). Purple lines are randomly selected single cell responses, black line is average well response. Graph is representative of three independent experiments conducted in duplicate wells. **c** Peak cytosolic Ca^2+^ responses in differentiated ReNcell VM matrices following administration of 10 nM GSK1016790A or DMSO control. Bar graph shows mean ± SEM of the averages of three independent experiments conducted in duplicate wells, data points are single cell values for all cells recorded across all experiments. Statistics were performed on average data results, unpaired two-tailed t test, ns; not significant. **d** Representative images of Ca^2+^ signaling responses in co-cultures of MDA-MB-468 breast cancer cells (expressing GCaMP6m) and ReNcell VM neural matrices (expressing jrCaMP1b) following administration of TRPV4 activator (10 nM GSK1016790A, dashed line indicates point of addition). Arrows highlight examples of ReNcell VM cells with Ca^2+^ activity. Timer indicates minutes:seconds. **e,g** Ca^2+^ responses in co-cultured MDA-MB-468 breast cancer cells **(e)** or ReNcell VM neural matrix cells (**g**) following addition of DMSO control (i) or TRPV4 activator GSK1016790A (ii). Green or purple lines are randomly selected single cell responses, black lines are well average. **f,h** Peak cytosolic Ca^2+^ responses in co-cultured MDA-MB-468 breast cancer cells (**f**) or ReNcell VM neural matrix cells (**h**) following addition of TRPV4 activator GSK1016790A or DMSO control. Bar graph is mean ± SEM of the averages of three independent experiments conducted in duplicate wells, data points are single cell values for all cells recorded across all experiments. Individual cells with values above 1.4 were set to 1.4 in fig **h** for plot scaling. Statistics were performed on average data results, unpaired two-tailed t test * *p* < 0.05.

Breast cancer and neural cell co-cultures expressing spectrally distinct Ca^2+^ sensors were exposed to either the TRPV4 activator GSK1016790A or the vehicle DMSO control and Ca^2+^ signaling was assessed via automated epi-fluorescence microscopy (Fig 2d, Supplementary Video 1). Spontaneous Ca^2+^ activity is a feature of both differentiated ReNcell VM neural matrices and MDA-MB-468 breast cancer cells in monoculture^41,52^. In vehicle DMSO control treated co-cultures, asynchronous spontaneous Ca^2+^ oscillations were observed in both cell populations (Fig 2*e i*, Fig 2*g i*, Supplementary Video 1). As expected, addition of the TRPV4 activator GSK1016790A to co-cultures rapidly induced a rapid and sustained influx of Ca^2+^ in MDA-MB-468 cells and significantly increased peak cytosolic free Ca^2+^ levels (Fig 2d, Fig 2e *ii*, Fig 2f, Supplementary Video 1). In contrast to monocultured ReNcell VM, co-cultured neural matrices had increased Ca^2+^ activity following administration of the TRPV4 activator (Fig 2d, Fig 2g *ii*, Supplementary Video 1). Despite a lack of endogenous TRPV4 expression, the peak Ca^2+^ level of ReNcell VM neural matrices in co-culture with TRPV4 expressing MDA-MB-468 breast cancer cells was significantly higher following administration of the TRPV4 activator GSK1016790A, in comparison to the addition of the vehicle DMSO control (Fig 2h). Responses of ReNcell VM in TRPV4 activator treated co-cultures were not uniform, however, and some neural cells appeared to have greater Ca^2+^ responses than others, particularly those close to breast cancer cells undergoing sustained increases in cytosolic Ca^2+^ (Fig 2d, arrows). To better elucidate the nature of Ca^2+^ signal changes in neural cells in co-culture with breast cancer cells, analysis of spatiotemporal relationships in these co-cultures was conducted.

### Ca^2+^ activity of neural cells is influenced by their proximity to stimulated breast cancer cells

To further define the nature of the observed Ca^2+^ responses, we turned our attention to analyzing our high content imaging data to identify characteristics of Ca^2+^ signal changes in the ReNcell VM matrices^53,54^. As discussed above, qualitative inspection revealed that Ca^2+^ signal increases in neural cells appeared more pronounced in regions adjacent to breast cancer cells exhibiting sustained Ca^2+^ elevations in response to GSK1016790A. (Supplementary video 1). To define spatial relationships within the co-cultures, we created a custom analysis approach utilizing Euclidean distance maps (Fig S2). Using this method, it was possible to determine the relative distances between all cells within the co-culture. Stratifying the neural cells in the co-culture as either proximate or distant to breast cancer cells (Fig 3a) started to reveal distinct responses in these groups (Fig 3b). In co-cultures treated with TRPV4 activator GSK1016790A, ReNcell VM cells proximate to breast cancer cells exhibited a greater increase in Ca^2+^ signal activity in contrast to neural cells more distant to breast cancer cells (Fig 3c *i, ii*). This enhanced activity was evident in both an increase in Ca^2+^ oscillation frequency and higher peak Ca^2+^ levels. Analysis of the average number of oscillations over time found that while proximate and distant ReNcell VM cells initially had similar rates of oscillations, only cells proximate to breast cancer had a trend of increased Ca^2+^ oscillations after selective breast cancer cell activation (Fig 3d *i*, *ii*). The trend of an increased number of Ca^2+^ oscillations was not immediately evident after cancer cell activation, but instead occurred only after 100 s post stimulation. This delay may reflect a requirement for significant Ca^2+^ transfer from cancer cells into neighboring neural cells requiring a sustained increase in breast cancer Ca^2+^ and/or the local sustained accumulation of paracrine signaling factors. The delay is also evidence that GSK1016790A did not directly activate ReNcell VM neural cells through mechanisms such as induced TRPV4 expression as a consequence of co-culture. ReNcell VM neural cells proximate to activated MDA-MB-468 breast cancer cells also had a significantly higher increase in peak Ca^2+^ compared to those that were more distant (Fig 3e).

**Fig 3:**
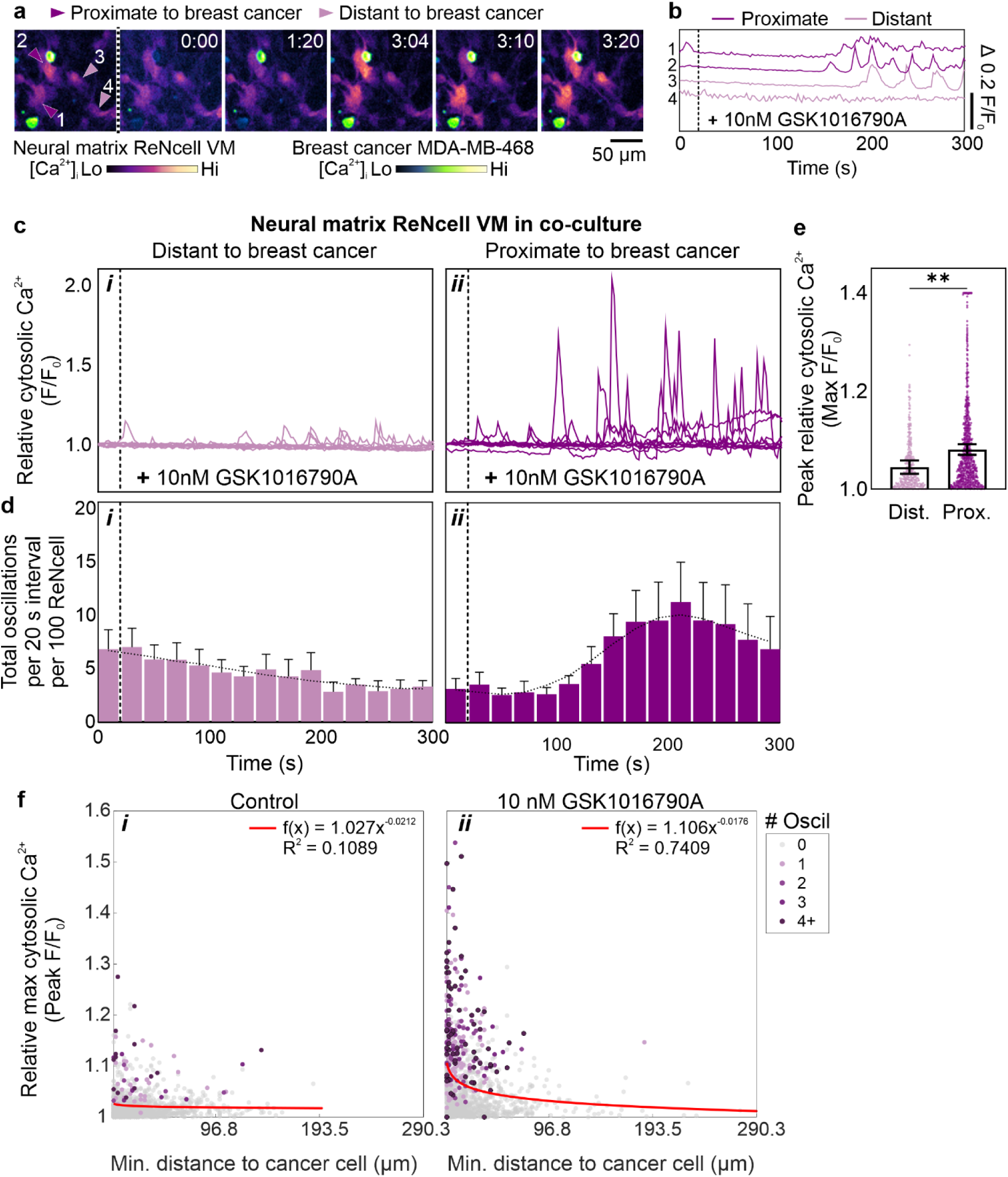
Communication of a Ca^2+^ signal from breast cancer to brain microenvironment is spatially dependent. **a** Representative time-lapse images of co-cultured MDA-MB-468 breast cancer and ReNcell VM neural matrix cells following addition of TRPV4 activator over 10 min (10 nM GSK1016790A, addition indicated by dashed line). Arrows indicate ReNcell VM classified as proximate (dark purple) or distant (light purple) to breast cancer cells. Images representative of three independent experiments. **b** Ca^2+^ traces of ReNcell VM cells depicted in **a**. **c** Ca^2+^ responses in co-cultured ReNcell VM neural cells either distant (***i***) or proximate (***ii***) to MDA-MB-468 breast cancer cells responding to TRPV4 activation in co-culture. Traces are 10 randomly selected single cell responses. **d** Peak cytosolic Ca^2+^ levels of ReNcell VM cells either distant or proximate to MDA-MD-468 breast cancer cells in co-culture. Bar graphs are mean ± SEM of the averages of three independent experiments conducted in duplicate wells, data points are single cell values for all cells recorded across all experiments. Single cell values above 1.4 were set to 1.4. Statistics were performed on average data results, unpaired two-tailed t test ** *p* < 0.05. **e** Histogram of oscillations over time in ReNcell VM either distant (***i***) or proximate (***ii***) to MDA-MB-468 breast cancer cells responding to TRPV4 activation (10 nM GSK1016790A, dotted vertical line represents time of addition) in co-culture. Bars are mean + SEM, curved line is fitted trend. Data are from three independent experiments conducted in duplicate wells. **f** Scatter plot of Ca^2+^ responses in ReNcell VM neural cells in co-culture with MDA-MB-468 breast cancer cells, following addition of DMSO control (***i***) or TRPV4 activator 10 nM GSK1016790A (***ii***). Dots are all single cell data from three independent experiments conducted in duplicate wells, red line is power regression curve fit to data. Color of dots represents number of oscillations detected during imaging period.

Fig 3f depicts the relationship between peak Ca^2+^ level, distance to nearest breast cancer cell, and total oscillations for each ReNcell VM in co-culture, following treatment with the TRPV4 activator or vehicle DMSO control (Fig 3f *i*, *ii*). A modest relationship between proximity and Ca^2+^ activity appeared present in the vehicle DMSO control group (Fig 3f *i*) which is consistent with the spontaneous Ca^2+^ oscillations that are a feature of MDA-MB-468 breast cancer cells in monoculture and maintained in our co-cultures with neural cells (Fig 2e *i*). However, this proximity relationship was much more pronounced in the group in which Ca^2+^ influx was selectively stimulated in breast cancer cells using GSK1016790A (Fig 3f *ii*). These results provide evidence that Ca²⁺ signaling from MDA-MB-468 breast cancer cells to ReNcell VM neural cells was spatially constrained and suggest that Ca²⁺ signal transmission may occurs preferentially at the tumor–brain microenvironment interface.

We believe that these results are the first simultaneous assessment of Ca^2+^ dynamics in neural and breast cancer cells with selective activation of cancer cells. Previous studies have suggested the possibility of such communication. For example, Zeng *et al.* found that expression of the Ca^2+^ permeable ion channel N-methyl-D-aspartate receptor (NMDAR) enhanced the survival of MDA-MB-231 breast cancer cells within the brain^29^. The authors concluded that NMDAR activation was dependent on signaling from the brain milieu, however, this conclusion was drawn from fixed samples and live cell Ca^2+^ assays of cancer cell lines in monoculture. More, recent work by Padmanaban *et al.* and Sanchez-Aguilera *et al.* have sought to explore Ca^2+^ communication between breast cancer and neuronal cells in the primary and metastatic environments, respectively^30,31^. Utilizing a co-culture model of breast cancer cell line 67NR and primary dorsal root ganglia (DRG) cells, Padmanaban *et al.* demonstrated that spontaneous Ca^2+^ events increased in DRG cells in co-culture in comparison to DRG cells in monoculture^31^. These Ca^2+^ events resulted in increased release of substance P from the DRG cells, which in turn promoted breast cancer growth and invasion. This work indicated that Ca^2+^ communication between breast cancer cells and neurons in the peripheral and primary tumor site settings may serve as a potential vulnerability for therapeutic targeting. However, the consequences of selective activation of breast cancer cells or neural cells on the Ca^2+^ dynamics of the other population was not assessed. In the metastatic context, Sanchez-Aguilera *et al.* conducted ex vivo bioluminescence Ca^2+^ imaging of murine brains injected with either melanoma, breast, or lung cancer cells^30^. However, the methodology employed by this study did not assess temporal or single cell Ca^2+^ dynamics. As both tumor and brain microenvironment expressed a bioluminescent Ca^2+^ sensor, it was not possible to distinguish Ca^2+^ activity in each discrete population. Our results now provide critical evidence to support the conclusion that cancer cells can remodel Ca^2+^ signaling in neural cells in a model of the brain metastatic site.

Intercellular communication via Ca^2+^ signaling can occur through a variety of mechanisms, including direct transfer of ions or signaling molecules, or paracrine secretion of signaling factors^55–58^. The spatially dependent Ca^2+^ signaling observed in this study suggests that Ca^2+^ communication is local. Such local Ca^2+^ communication has been observed in metastatic models of breast cancer in other metastatic sites. In a co-culture model of MCF-7 breast cancer cells and osteogenic cells, connexin-43 dependent gap junctions allow the transfer of Ca^2+^ from the bone microenvironment to cancer cells, enhancing survival of breast cancer at this site^59^. Alternately, induction of Ca^2+^ mobilisation in astrocytes can cause exocytosis of adenosine triphosphate and could have contributed to response of ReNcell VM cells not in direct contact with breast cancer cells^60,61^. A Ca^2+^ signal could also be propagated through the elongated cell bodies of the ReNcell VM cells, further cascading the Ca^2+^ signal even in neural cells far away from stimulated breast cancer cells. The exact mechanism of Ca^2+^ communication between breast cancer and the brain microenvironment has not been fully established in any studies exploring the importance of Ca^2+^ signaling in brain metastases^29,30^. Further work defining these exact mechanisms may provide fruitful opportunities for new therapeutic intervention approaches. Given the distinct Ca²⁺ signaling dynamics between breast cancer and neural cells, we next investigated a potential consequence of this spatially dependent phenomenon, by assessing the expression of a Ca²⁺-responsive gene using RNA fluorescence in situ hybridization (RNA-FISH).

### ReNcell VM neural microenvironment cells undergo spatially related expression remodeling when co-cultured with breast cancer cells

Expression remodeling adaptations in the brain microenvironment are a known consequence of metastatic invasion^33,62,63^. To explore whether this was also observable in our model, we conducted RNA-FISH, which allows quantification of gene expression levels while retaining spatial information^64,65^. SRY-Box Transcription Factor 2 (SOX2) is a transcription factor linked to neural differentiation and key cancer processes^66^. Expression of SOX2 is Ca^2+^ signal dependent in chick embryos, and remodeled in brain stroma near metastatic lesions^62,67^. Previous studies have found that ReNcell VM express SOX2, while MDA-MD-468 cells have undetectable expression of this target^40,68,69^. To identify the breast cancer cells within the co-cultures, we also stained for expression of TRPV4, a gene exclusively expressed by MDA-MD-468 cells in this co-culture model of breast cancer brain metastasis (Fig 2a)^51^. SOX2 expression in the ReNcell VM matrix was correlated with spatial proximity of neural cells to MDA-MB-468 breast cancer cells (Fig 4a, b). ReNcell VM cells proximate to breast cancer had higher expression levels of SOX2 in comparison to ReNcell VM distant to breast cancer cells (Fig 4b). This result is consistent with the ex-vivo model of Sanchez-Aguilera *et al.* which demonstrated increased Ca^2+^ dependent secretion of substance P in DRG cells in co-culture with breast cancer cells in the absence of activation of breast cancer cells^30^.

**Fig. 4:**
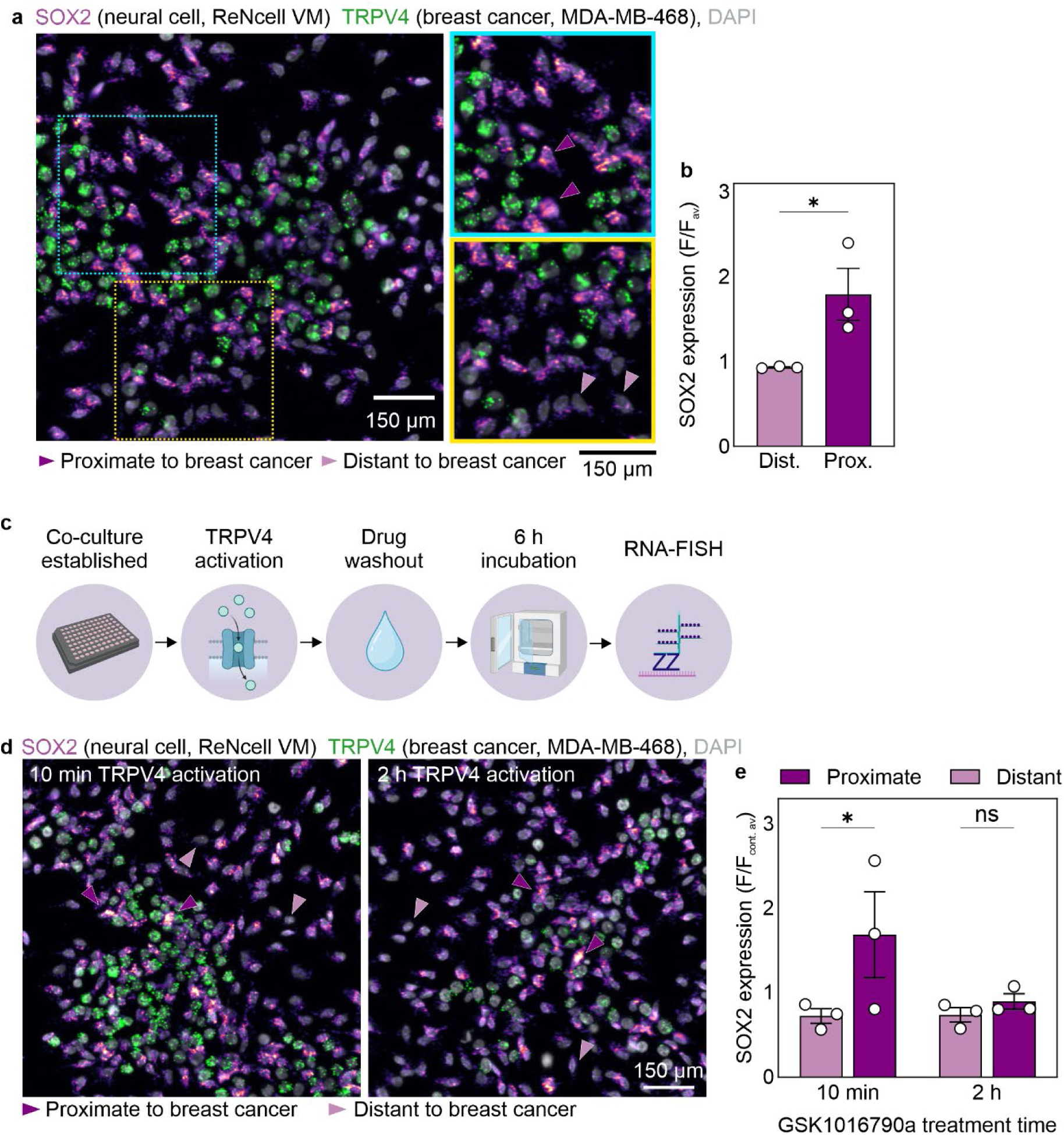
Co-culturing of breast cancer and neural matrices results in spatially dependent expression remodeling in the brain microenvironment. **a** Representative images of RNA-FISH staining for TRPV4 and SOX2 RNA in MDA-MB-468 breast cancer and ReNcell VM neural matrix co-cultures. Arrows indicate ReNcell VM cells either distant (light purple) or proximate (dark purple) to breast cancer cells. Images representative of three independent experiments conducted in duplicate. **b** SOX2 expression in ReNcell VM cells either distant or proximate to breast cancer cells in co-culture. Bars are mean ± SEM, dots are experiment average. Data are from three independent experiments conducted in duplicate wells, significance was assessed via unpaired t test, * *p* < 0.05 **c** Flow-chart of experimental process for data depicted in **d** and **e**. **d** Representative images of RNA-FISH staining for TRPV4 and SOX2 RNA in co-cultures treated with TRPV4 activator GSK1016790A for 10 min or 2 h. Arrows indicate ReNcell VM cells either distant (light purple) or proximate (dark purple) to breast cancer cells. Images representative of three independent experiments conducted in duplicate. **e** SOX2 expression in ReNcell VM either distant or proximate to breast cancer cells following 10 min or 2 h of TRPV4 activation. Bars are mean ± SEM, dots are experiment average. Data are from three independent experiments conducted in duplicate wells, significance was assessed via two-way ANOVA, * *p* < 0.05.

To explore the possibility that stimulated Ca^2+^ communication could influence SOX2 expression in the ReNcell VM matrices, we conducted RNA-FISH in co-cultures following treatment with the TRPV4 activator GSK1016790A (Fig 4 c). Analysis of SOX2 expression in this context found that following 10 min of TRPV4 activation, proximate ReNcell VM had higher expression than ReNcell VM that were distant to breast cancer (Fig 4d,e). However, more extended treatment with the TRPV4 activator resulted in the loss of this correlation. This may reflect a sustained activation pathway across the entire neural cell monolayer. Taken together, these SOX2 expression studies are consistent with previous reports that proximity to metastatic breast cancer cells results in expression remodeling of the brain microenvironment, and further reinforce the importance of the spatial relationship between breast cancer and neural cells to induce this remodelling^31,62,63^. Furthermore, there was an absence of expression of TRPV4 in ReNcell VM within co-culture (Fig 4a&d). This demonstrates that the Ca^2+^ responses observed in our neural matrices following TRPV4 activation were due to communication of a Ca^2+^ signal, and not due to TRPV4 expression being induced in neural cells as a result of co-culturing with breast cancer cells.

Our work describes a high-throughput, human cell model that was well suited for exploring the Ca^2+^ signaling dynamics between breast cancer cells and the brain microenvironment at a single cell level with high temporal resolution. Future adaption of this model could potentially be utilized to evaluate emerging evidence in animal models that some brain metastases have identifiable subtype-specific impacts on neural activity^30^. A caveat in our current model is the inability to distinguish between neural cell types while conducting live Ca^2+^ imaging. A potential means to circumvent this in the future could be to drive expression of spectrally distinct genetically encoded Ca^2+^ indicators downstream of cell type specific promoters, such as GFAP or β3-tubulin. Such an approach would represent a natural progression from the single neural cell analysis that was achieved in these studies to neural cell sub-types of single cells and allow future comparisons of neurons and glia in the context of breast cancer brain metastasis.

The results presented in this study provides the first simultaneous assessment of Ca^2+^ signaling in breast cancer cells and neural cells in a model of the brain metastatic microenvironment. Selective activation of Ca^2+^ influx in breast cancer cells is sufficient to induce a significant remodeling of Ca^2+^ signaling in proximal neural cells. Given breast cancer brain metastasis remains a pressing clinical challenge due to a lack of targeted therapeutic options^4,8,9^, ongoing efforts to understand and disrupt Ca^2+^ signal communication within the brain metastatic niche may offer new avenues for intervention.

## Methods

### Cell culture

ReNcell VM were purchased from Merck (Germany), MDA-MB-468 cells were obtained from The Brisbane Breast Bank, UQCCR, Australia. All cells were maintained in a humidified incubator at 37°C with 5% CO_2_, routinely screened for mycoplasma at the Translational Research Institute, Brisbane Australia, and authenticated via short tandem repeat (STR) profiling at QIMR Berghofer, Brisbane, Australia. ReNcell VM were cultured in the proliferative state on laminin coated (12 µg/mL, ThermoFisher Scientific, USA) plasticware with ReNcell Maintenance Medium (Merck) supplemented with epidermal growth factor (EGF, 20 ng/mL, Merck) and basic fibroblast growth factor (bFGF, 20 ng/mL, Merck). MDA-MB-468 cells were cultured as previously described^70^.

To generate co-cultures, proliferative ReNcell VM were plated on laminin coated black walled 96-well plates (Falcon, InVitro Technologies, Australia) and maintained in ReNcell Maintenance Medium for 48 h prior to withdrawal of mitogens to induce differentiation. ReNcell VM monolayers were differentiated for 96 h prior to addition of MDA-MB-468 cells. Co-cultures were maintained in ReNcell Maintenance Medium supplemented with 0.5% fetal bovine serum (FBS, Moregate Biotech, New Zealand) for 7 d prior to further experiments. Culture medium was refreshed every 48 h during this time. Co-cultures were generated using either wild-type ReNcell VM and MDA-MB-468 cell lines, or the same cell lines stably expressing genetically encoded Ca^2+^ indicators (generation of which is outlined below).

### Lentiviral expression of genetically encoded Ca^2+^ indicators

MDA-MB-468 cells expressing genetically encoded Ca^2+^ indicator GCaMP6m were developed as previously described^71^. ReNcell VM expressing genetically encoded Ca^2+^ indicator jrCaMP1b (pGP-CMV-NES-jRCaMP1b, Addgene) were developed via cloning the jrCaMP1b sequence into lentiviral vector pCDH-EF1-FHC (Addgene) via BamHI and EcoRI restriction sites. Lentiviral vectors pCMV-VSV-G, pCMV-dR8.2 dvpr (Addgene) and pCDH-EF1-FHC-jrCaMP1b were transfected into HEK293T cells using Lipofectamine 3000 (ThermoFisher Scientific). The day after transfection, HEK293T media was replaced with ReNcell Maintenance Medium supplemented with growth factors as described above for 24 h prior to supernatant removal. Virus-containing supernatant was cleared by centrifugation and stored at −80°C for a least 1 h. Proliferative ReNcell VM were transduced with virus preparation (4.5 mL viral supernatant, 5.5 mL of ReNcell Maintenance Medium supplemented with a final concentration of 20 ng/mL for both EGF and bFGF, and 40 µg polybrene, Merck). After 24 h, the medium was replaced with fresh ReNcell Maintenance Medium supplemented with growth factors. Puromycin (0.5 µg/mL, Sigma-Aldrich, USA) selection was commenced 48 h post viral infection.

### RT-qPCR

RNA was isolated via RNeasy Plus Mini Kit (Qiagen). Samples were reverse transcribed into cDNA using the Omniscript Reverse Transcription Kit (Qiagen), with RNase inhibitor (RNasin® Ribonuclease Inhibitor, Promega) and random primers (Promega) purchased separately. cDNA was amplified via Taqman Fast Universal PCR Master Mix (Life Technologies, USA) and Taqman gene expression assays in a StepOnePlus Real-Time PCR System (Life Technologies). Relative mRNA levels were calculated via comparative C_T_ method, normalised to endogenous controls: 18S for MDA-MB-468 cells, or glyceraldehyde-3-phosphate dehydrogenase (GAPDH) for ReNcell VM. Taqman real-time assays used in this study: 18s 4319413E, GAPDH Hs99999905_m1, GFAP Hs00909233_m1, MAP2 Hs00258900_m1, TRPV4 Hs00219765_m1.

### Immunofluorescence

ReNcell VM were plated in 96-well black walled plates (Falcon, InVitro Technologies, Australia) and differentiated for 96 h prior to fixing in 4% paraformaldehyde (ThermoFisher).Following fixing, cultures were incubated for 5 min in a permeabilisation buffer (0.5% Triton-X 100 (Sigma-Aldrich), 0.1% Bovine Serum Albumin (BSA, Sigma-Aldrich) in PBS), and then a blocking buffer (5% BSA) for 30 minutes at room temperature. Primary-conjugated antibodies were diluted in blocking buffer and incubated overnight at 4°C. Antibodies used in this study were: Anti-GFAP Cy3 primary conjugate (1:200, MAB3402C3, Merck), and Anti-tubulin βIII Alexa-Fluor 488 (1:50, clone 2610-TB3, 41-4510-80, ThermoFisher). Following primary antibody incubation, samples were stained for 10 min with nuclear counterstain DAPI (Invitrogen, USA). Cultures were imaged via automated epi-fluorescence microscopy in an ImageXpress Micro (Molecular Devices, USA) with 10x objective and following filter cubes: GFP (excitation: 472/30 nm; emission: 520/35 nm), Cy3 (excitation: 531/40 nm; emission: 593/40 nm) and DAPI (excitation: 377/50 nm; emission: 477/60 nm). The Multi Wavelength Cell Scoring Tool in the MetaXpress software (Version 7.0, Molecular Devices) was used to characterize the proportions of each cell type in the neural cultures.

### Brightfield imaging

Co-cultures were imaged via a JuLI Stage (NanoEnTek, Korea), located within a humidified incubator at 37°C and 5% CO_2_.

### High-content imaging of intracellular Ca^2+^ via automated epifluorescence microscopy

ReNcell VM and MDA-MB-468 monoculture or co-cultures were prepared as described above. Immediately prior to imaging, plates had media removed and replaced with Fluorobrite DMEM Media (ThermoFisher) supplemented with 4 mM L-glutamine (ThermoFisher), 25 mM HEPES (ThermoFisher) and 0.5% FBS. Cultures were imaged at 37°C on an ImageXpress Micro using the GFP and Cy3 filter cubes (Ex/Em information provided above). Monoculture imaging was conducted on 10x objective with no binning, image frequency of 1 s, duration of 151 s, and carbachol (Merck) addition after 120 s. Co-culture images were taken on the 10x objective, with 3 x 3 binning, an imaging frequency of 2 s, duration of 300 s and DMSO (Sigma-Aldrich) or GSK1016790A (Sigma-Aldrich) addition after 20 s.

### RNA fluorescent in situ hybridisation (RNA-FISH) and imaging

RNA-FISH was performed on co-cultures of differentiated ReNcell VM and MDA-MB-468 cells in 96-well plates using the RNAScopeTM Multiplex Fluorescent V2 Assay Kit (ACD Inc, Newark, CA) according to the manufacturer’s protocol. Briefly, cells were washed in PBS and fixed using 10% neutral buffered formalin for 30 min at room temperature. Cells were dehydrated using 50%, 70% and 100% ethanol and stored at −20°C. On the day of hybridization, cells were rehydrated then treated with hydrogen peroxide for 10 min, washed with dH2O and then digested using Protease III for 10 min at room temperature. Cells were then washed and hybridized with target probes, TRPV4-C1 (Cat. #452221) and SOX2-C2 (Cat. #400871C2) at 40°C for 2 h in a humidified environment. Cells were then washed and incubated with 5X SSC buffer overnight at room temperature. Signal amplification was performed by incubating with AMP1, AMP2 and AMP3 probes and HRP signals in C1 and C2 channels were developed by subsequent incubations with HRP-C1, then Opal-520 fluorophore dye (Akoya Biosciences, Marlborough, MA) and HRP-C2, then Opal-570. Cell nuclei were stained with DAPI. Images were acquired using an automated epifluorescence microscope (ImageXpress Micro, Molecular Devices) with a 20X objective and GFP (excitation: 472/30 nm; emission: 520/35 nm), Cy3 (excitation: 531/40 nm; emission: 593/40 nm) and DAPI (excitation: 377/50 nm; emission: 477/60 nm) filter cubes.

### Image quantitation

Image quantification was conducted via MetaXpress software using a custom designed analysis journal based on a previously established protocol to identify single cells and record fluorescence values over time^54^. Briefly, a segmentation mask for each wavelength was generated, and potential bubbles and cells at the edge of imaging field were identified within the mask and removed. Following this, relative distances of cells within co-cultures were determined. Sequentially for each MDA-MB-468 cell, a Euclidean distance map was generated which created an intensity gradient originating from the perimeter of the cell undergoing analysis. Using this, relative distances from the perimeter of the breast cancer cell undergoing analysis to each cell in the segmentation mask for the ReNcell VM could be calculated. Supplementary Fig S2 provides a visual depiction of this process.

Images were visualized and prepared via ImageJ (version 2.3.0, National Institutes of Health) using the BIOP LUT^72^.

### Data analysis

Data was analyzed via MATLAB software (R2021b, MathWorks, USA). For Ca^2+^ imaging data, fluorescence values were normalized against initial recorded value (F/F_0_), for RNA-FISH experiments, individual fluorescence values were normalized against the average fluorescence of control wells for each experiment. Ca^2+^ oscillations were identified using the MATLAB findpeaks function, with a minimum peak prominence set at 0.3 F/F_0_. In Fig 3 and 4, ReNcell VM were classified as distant to breast cancer cell if the distance between the perimeter of the ReNcell VM and the nearest breast cancer cell was greater than average cell width of the MDA-MB-468 population. ReNcell VM at distances less than this were classified as proximate to breast cancer. Power regression line fit and R^2^ values were determined using the MATLAB curve fitting tool.

### Statistical analysis

Statistical analysis was performed using GraphPad Prism (Version 10.1.0). Statistical tests are outlined in figure legends.

## Data and code availability

The datasets generated and analyzed via custom MetaXpress and/or MATLAB algorithms from this study are available from the corresponding author on reasonable request.

## Acknowledgements

This work was supported by the Office of the Assistant Secretary of Defense for Health Affairs, through the Breast Cancer Research Program, under Award No. W81XWH-17-1-0064. Opinions, interpretations, conclusions and recommendations are those of the author and are not necessarily endorsed by the Department of Defense. Figure 4 in this manuscript utilized BioRender icons. Chalmers, S. (2025) https://BioRender.com/ma2trko

## Author Contributions

All authors contributed to the study conception and design. S.B.C, A.H.L.B, and M.R performed the experiments. S.B.C and F.S analyzed the Ca^2+^ signaling data, S.B.C performed all other data analyses. The first draft of the manuscript was written by S.B.C and G.R.M. All authors read, edited and approved the final manuscript.

## Materials and correspondence

Correspondence and material requests should be sent to G. R. Monteith

## Figures

**Fig S1.**
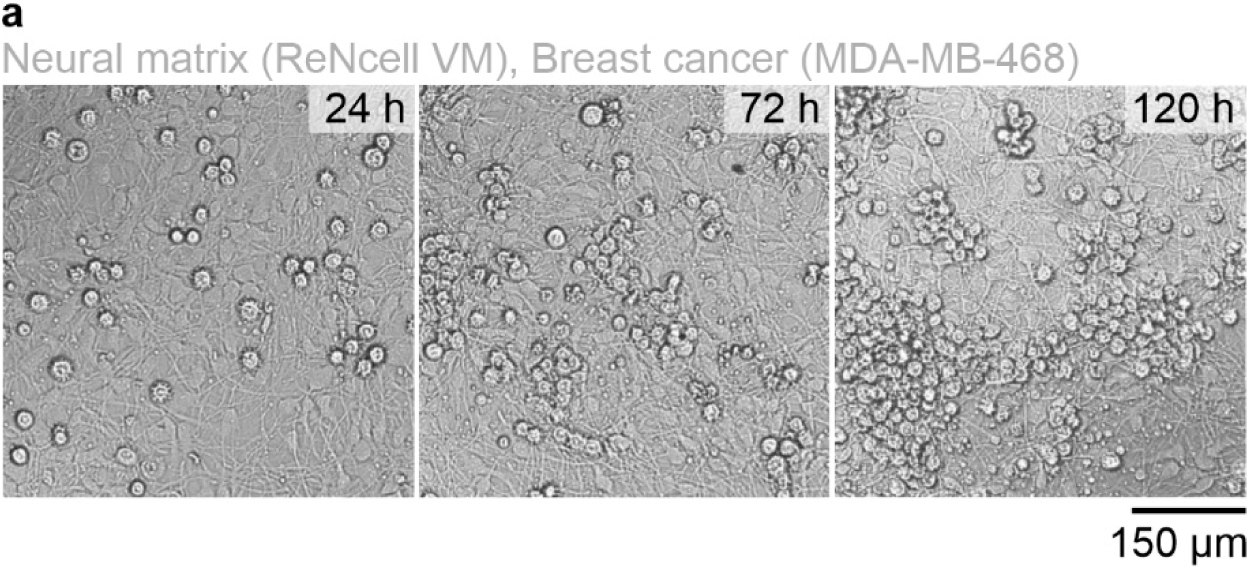
Human neural progenitor cell line, ReNcell VM sustains breast cancer growth. **a** Raw data of brightfield images from Fig 1a, depicting differentiated ReNcell VM neural matrices (spindle shaped cells) supporting the growth of breast cancer cell line MDA-MB-468 (cobblestone shaped cells) over time.

**Fig S2.**
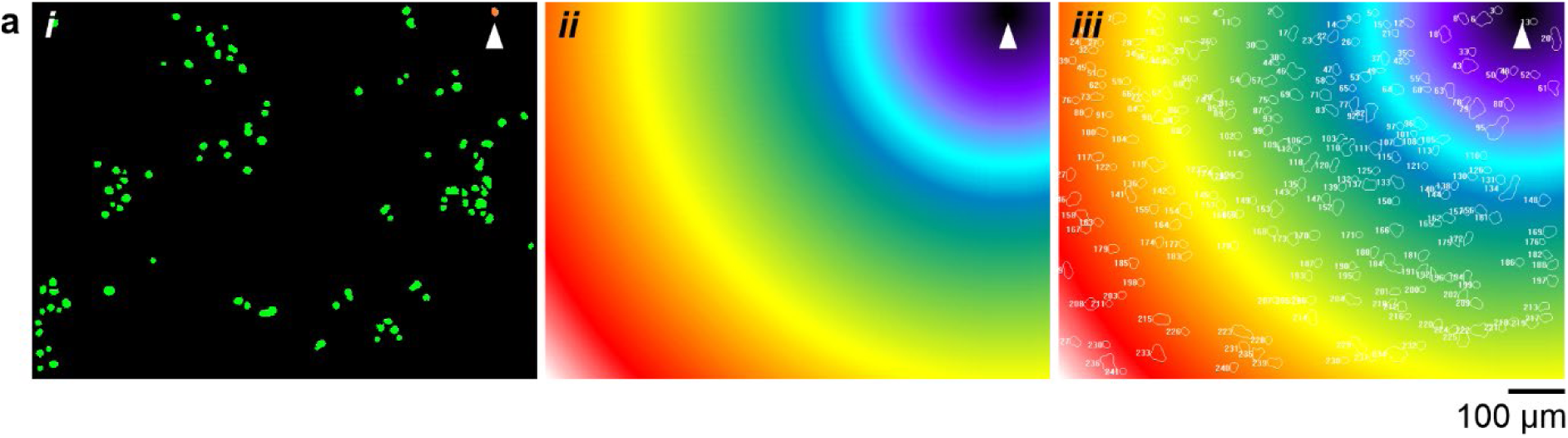
Automated identification of relative cell distances in co-culture utilizing a Euclidean distance map approach. **a *i*** Segmentation mask for MDA-MB-468 breast cancer cells in co-culture with ReNcell VM neural matrix. Arrow indicates individual MDA-MB-468 breast cancer cell undergoing distance analysis. ***ii*** Euclidean distance map generated for cell indicated in panel ***i*.** The Euclidean distance map creates an intensity gradient originating from the cell perimeter which provides information on relative distances within the image as each additional pixel from the cell perimeter increases the intensity by 1. ***iii*** Transposition of the segmentation mask for ReNcell VM cells in co-culture onto the Euclidean distance map allows for measurement and recording of relative distances between MDA-MB-468 in question and each ReNcell VM in the imaging field. This process is repeated for each MDA-MB-468 cell in the image.

**Supplementary video 1. Treatment of breast cancer and neural co-cultures with TRPV4 activator GSK1016790A or control.** Co-cultures of MDA-MB-468 breast cancer (green cells) and ReNcell VM neural matrices (purple cells) with DMSO control (left video) or TRPV4 activator GSK1016790A (10 nM, right video). Timer indicates minutes:seconds. Video is representative of three independent experiments conducted in duplicate wells.

